# Measuring the metabolic evolution of glioblastoma throughout tumor development, regression, and recurrence with hyperpolarized magnetic resonance

**DOI:** 10.1101/2021.06.10.447987

**Authors:** Travis C. Salzillo, Vimbai Mawoneke, Joseph Weygand, Akaanksh Shetty, Joy Gumin, Niki M. Zacharias, Seth T. Gammon, David Piwnica-Worms, Gregory N. Fuller, Christopher J. Logothetis, Frederick F. Lang, Pratip K. Bhattacharya

## Abstract

Rapid diagnosis and therapeutic monitoring of aggressive diseases such as glioblastoma can improve patient survival by providing physicians the time to optimally deliver treatment. This research tested whether metabolic imaging with hyperpolarized MRI could detect changes in tumor progression faster than conventional anatomic MRI in patient-derived glioblastoma murine models. To capture the dynamic nature of cancer metabolism, hyperpolarized MRI, NMR spectroscopy, and immunohistochemistry were performed at several time-points during tumor development, regression, and recurrence. Hyperpolarized MRI detected significant changes of metabolism throughout tumor progression whereas conventional MRI was less sensitive. This was accompanied by aberrations in amino acid and phospholipid lipid metabolism and MCT1 expression. Hyperpolarized MRI can help address clinical challenges such as identifying malignant disease prior to aggressive growth, differentiating pseudoprogression from true progression, and predicting relapse. The individual evolution of these metabolic assays as well as their correlations with one another provides context for further academic research.

## Introduction

In 2020, an estimated 87,240 new cases of primary brain and other central nervous system (CNS) tumors were diagnosed in the United States with 25,800 expected to be malignant (1). WHO grade IV glioblastoma (GBM) is the most common malignant brain tumor in adults representing 48.3% of the cases as well as the most deadly with 5-year survival rates of merely 6.8%. Despite standard-of-care treatment with surgery, radiotherapy, and temozolomide chemotherapy, prevailing therapies remain palliative, and tumors are almost always recurrent and lead to median survival times of approximately 15 months (2). Alternative therapies, both FDA-approved and experimental, have shown little to no improvement on overall survival, as factors such as relatively average mutational load, tumor heterogeneity, and molecular filtration by the blood-brain barrier challenge drug development for this disease (3). This failure to make major inroads points to the need for alternative approaches in the management of this disease.

Imaging has been used to inform anatomic-based intervention by determining the extent of involvement of the tumor, its proximity to functional regions, and monitoring recurrence. Advanced imaging techniques provide the opportunity to examine the cancer in alternate biologic domains that may determine therapeutic vulnerability in specific subsets. Such a strategy would complement the prevailing anatomic-based classifiers and could potentially address several clinical challenges when it comes to improving patient survival and optimizing drug discovery. These include the early detection and discrimination of malignant disease, rapid assessment of treatment efficacy (including distinguishing pseudo-progression from true progression), and prediction of tumor recurrence. Molecular imaging, especially by probing tumor metabolism, has shown great promise in augmenting conventional clinical imaging for the diagnosis, prognosis, and treatment monitoring of brain tumors (4, 5).

Hyperpolarized magnetic resonance (MR) is one such technique that has found success over the past several years for diagnosing a number of tumor types (6) and measuring their response to therapy (7, 8). Pyruvate is the most commonly used hyperpolarized substrate due to its relatively long signal enhancement lifetime and central role in tumor metabolism via the Warburg effect (9, 10). Compelling findings from the numerous preclinical studies with this substrate have justified the initiation of several clinical trials for the use of hyperpolarized pyruvate in multiple tumor sites including brain cancer, which has produced human data from healthy volunteers (11–13) and glioma patients (14–17). Given the current advances, there is a strong likelihood that hyperpolarized pyruvate MR will be adopted in the clinic with FDA approval for metabolic imaging of brain cancer, and protocols for integrating this technique into the clinical workflow is underway (18).

There is still much to be explored to determine the specific clinical implementations of hyperpolarized MR to produce significant impact in the treatment of brain tumors. Existing preclinical studies do a thorough job of elucidating differences in pyruvate utilization between tumors and healthy brain (19–23) or between untreated and treated tumors (24–27), but they only do so at one point during development or with one pre- and post-treatment measurement. Tumor metabolism is heterogeneous and evolves over the course of tumor development suggesting that the time-point of the measurement can significantly impact the results (28–30). Following treatment, metabolism is similarly dynamic as it responds to therapeutic insults before gaining resistance or re-growing. This complex metabolic trajectory cannot be captured with one post-treatment measurement alone. Therefore, we sought to implement serial hyperpolarized MRS measurements at multiple time-points over the course of tumor growth and treatment regimen to elucidate this metabolic evolution. *In vivo* hyperpolarized pyruvate-to-lactate conversion values were determined at multiple time-points throughout three stages of tumor progression (development, regression following radiotherapy and recurrence to the point of relapse). Additionally, the novel use of hyperpolarized MRS to assess brain tumor recurrence, which is an unfortunate and inevitable reality for most GBM patients, could improve patient survival by informing physicians when additional treatment is necessary before the tumor aggressively regrows (31, 32).

The purpose of this study was to compare hyperpolarized pyruvate-to-lactate conversion values from the serial hyperpolarized MRS experiments with tumor volume changes that were acquired with anatomic MRI during each stage of tumor progression (Fig. 1). This was done to demonstrate the value of adding hyperpolarized MR to conventional imaging protocols by addressing several challenges commonly encountered in the clinical setting. Furthermore, we sought to form a more complete picture of the metabolic events occurring during tumor progression by performing metabolomics with nuclear magnetic resonance (NMR) spectroscopy and protein expression assays with immunohistochemistry (IHC) on *ex vivo* samples at multiple time-points and investigating the interrelationships of each of these measurements over the entire course of tumor lifetime.

**Figure 1:**
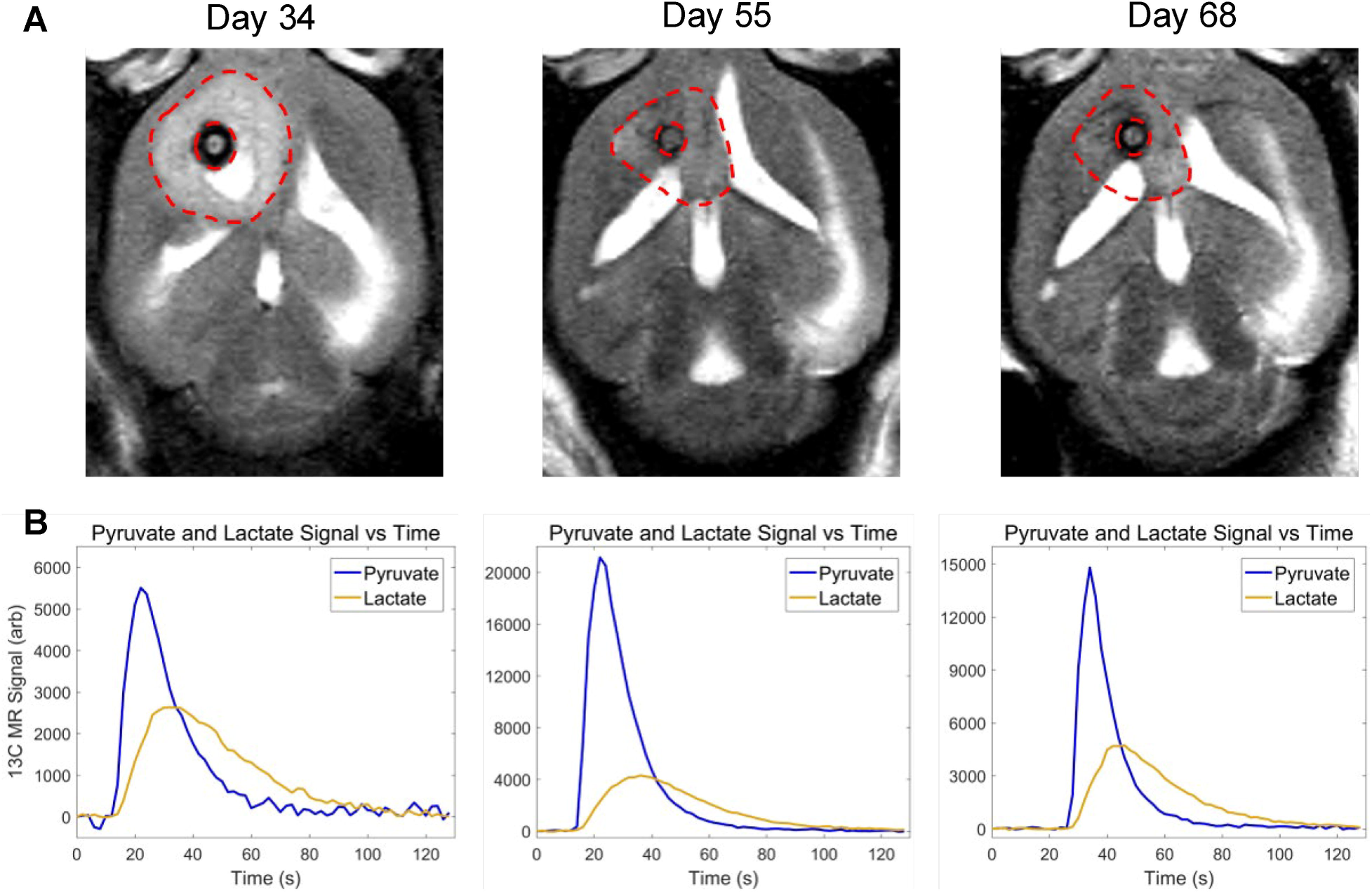
Anatomic and metabolic imaging of tumor-bearing mice over time. Tumor volume **(A)**, imaged with T2-weighted MRI, and pyruvate-to-lactate conversion **(B)**, measured with hyperpolarized MRS, is displayed at the end of tumor development (Day 34), end of tumor regression (Day 55), and at the point of relapse (Day 68) in the same mouse.

## STAR Methods

### RESOURCE AVAILABILITY

#### Lead Contact

Further information and requests for resources and reagents should be directed to and will be fulfilled by the Lead Contact, Pratip Bhattacharya, Ph.D.

#### Materials Availability

This study did not generate new unique reagents.

#### Data and Code Availability

The dataset containing all tumor volume, nLac, metabolite pool size, and protein expression values generated during this study as well as the MATLAB code used to calculate nLac from raw hyperpolarized ^13^C spectral data are available at http://dx.doi.org/10.17632/mf5f93t3kn.1

### EXPERIMENTAL MODEL AND SUBJECT DETAILS

#### Cell Lines

GSC 8-11: Glioma sphere-forming cells (GSC) 8-11 were isolated from surgical sample from a female patient which have been reported in the literature (Mansouri et al., 2016). As reported in the literature (Singh et al., 2004; Wei et al., 2010), cells were grown in Neurosphere Media containing DMEM/F12 (Corning, Corning, NY, USA) with B27 (x1, Thermo Fisher Scientific, Waltham, MA, USA), bFGF (20 ng/ml, Millipore Sigma, St. Louis, MO, USA), and EGF (20 ng/ml, Millipore Sigma, St. Louis, MO, USA) at a temperature of 37° C. These cells were authenticated by the MDACC Cell Authentication Core (https://www.mdanderson.org/research/research-resources/core-facilities/cytogenetics-and-cell-authentication-core.html).

#### Animals

<u>Mice:</u> Five-week-old athymic nude mice (Experimental Radiation Oncology, MDACC, Houston, TX, USA) were used for *in vivo* studies. Mice were housed together in a sterilized facility with 5 to a cage, so only females were used as they are less-prone to fighting with each other. Mice received standard feed and water and were inspected for health daily. A guide-screw system was used for injection to allow for consistent placement of intracranial xenografts as described in literature (Lal et al., 2000). A 2.6 mm long guide screw with a 0.5 mm channel bored through its center was drilled into the skull directly over the caudate nucleus. After the mice had recovered for 2 weeks, 5 × 10^5^ GSC 8-11 cells were suspended in 3 µL of phosphate-buffered saline (PBS) and injected stereotactically through the bore of the guide screw over a period of five minutes. After injection, a stylet was placed in the bore of the screw to close the system and prevent tissue from growing inside. Control animals were prepared in the same manner except that PBS absent of GSCs was injected. All procedures were performed in accordance with regulations of the Institutional Animal Care and Use Committee (IACUC) of the University of Texas MD Anderson Cancer Center.

### METHOD DETAILS

#### Experimental Overview

Following the intracranial implantation of patient-derived glioma sphere-forming cells (GSC), the anatomic and metabolic properties of this GBM model were interrogated at several time-points during three stages of tumor progression: tumor development, regression following radiotherapy, and eventual recurrence. Mice were split into three cohorts: untreated tumor-bearing mice, treated tumor-bearing mice (2x5 Gy radiotherapy), and control mice. Anatomic growth and shrinkage were studied *in vivo* with T1-weighted, T2-weighted, and fluid-attenuated ^1^H MRI. Real-time conversion of injected pyruvate to lactate was measured in the tumor *in vivo* with hyperpolarized ^13^C MRS. *Ex vivo* metabolite pool sizes and protein expression were determined with NMR spectroscopy and IHC, respectively. All measurements were acquired within ±1 day of their nominal time-point.

#### Tumor Radiotherapy

Tumor-bearing mice in the treatment cohort underwent radiotherapy to attenuate tumor growth. On Days 25 and 27, the mice were imaged and treated with 5 Gy whole-brain irradiation on the X-RAD SmART small animal irradiator (Precision X-Ray, North Branford, Connecticut, USA). A cone-beam CT was acquired for treatment planning, and irradiation was executed with two opposing fields of 2.5 Gy each.

#### Anatomic Magnetic Resonance Imaging

Tumor anatomy was visualized with MRI on a Bruker 7 T pre-clinical scanner (Bruker Biospin MRI GmbH, Ettingen, Germany) every 3 days. T1-weighted (T1-w), T2-weighted (T2-w), and fluid attenuated inversion recovery (FLAIR) ^1^H pulse sequences were implemented. A 35 mm RF volume coil (Bruker) was used to acquire the images. The following pulse sequences and parameters were used: coronal, sagittal, and axial T2-w (FA = 90°, TE = 6.5 ms, TR = 1500 ms, BW = 75 kHz, Matrix = 256 x 192, FOV = 25 mm x 25 mm (axial 20 mm x 20 mm), NEX = 4, Slice Thickness = 0.75 mm, Slice Gap = 0.25 mm, RARE Factor = 12); axial T2-w FLAIR (FA = 90°, TE = 48 ms, TR = 10,000 ms, TI = 2000 ms, BW = 38 kHz, Matrix = 256 x 192, FOV = 20 mm x 20 mm, NEX = 3, Slice Thickness = 0.75 mm, Slice Gap = 0.25 mm, RARE Factor = 10, Fat Suppression on); axial T1-w (FA = 90°, TE = 57 ms, TR = 3000 ms, BW = 85 kHz, Matrix = 256 x 192, FOV = 20 mm x 20 mm, NEX = 3, Slice Thickness = 0.75 mm, Slice Gap = 0.25 mm, RARE Factor = 4, Fat Suppression on).

#### Tumor Volume Measurements

DICOM images from the T1-w, T2-w, and FLAIR pulse sequences were imported into 3D Slicer software (Fedorov et al., 2012). Tumors were segmented on each slice of the coronal, sagittal, and axial T2-w images. T1-w and FLAIR images were used to confirm tumor boundaries and differentiate tumor tissue from surrounding edema and cerebral spinal fluid. Volume in each plane of the T2-w images was calculated by multiplying the number of voxels within the tumor segmentation by the spatial dimensions of the voxels. The calculated volumes from each of the three T2-w imaging planes were averaged together to give the final volume measurement of the tumor. At least 5 tumor volumes were averaged at each time-point.

#### Hyperpolarized Sample Preparation

[1-^13^C]pyruvic acid (Millipore Sigma, St. Louis, MO, USA) was doped with Ox063 trityl radical (Oxford Instruments, Abingdon, UK) to 15 mM concentration. 20 µL of this solution was mixed with 0.4 µL of 50 mM Gd^3+^ (Bracco Diagnostics, Monroe Township, NJ, USA). This solution was placed in a DNP HyperSense (Oxford Instruments, Abingdon, UK) to polarize for approximately 1 hour under microwave irradiation at 94,100 GHz. An average polarization of 20,000 was achieved. Once the sample was prepared, it was rapidly heated and dissolved in 4 mL buffer comprised of 40 mM 2-Amino-2-(hydroxymethyl)-1,3-propanediol (TRIS; Millipore Sigma, St. Louis, MO, USA), 80 mM NaOH, 0.1 g/L ethylenediaminetetraacetic acid (EDTA; Millipore Sigma, St. Louis, MO, USA), and 50 mM NaCl. This solution had a final [1-^13^C]pyruvic acid concentration of 80 mM which was then injected intravenously through the tail vein of the mouse.

#### 13C Magnetic Resonance Spectroscopy

A 72 mm ^1^H volume coil (Bruker Biospin MRI GmbH, Ettingen, Germany) was used to acquire anatomic images for accurate region of interest (ROI) placement for spectroscopy. A ^13^C transmit/receive surface coil (ID: 35 mm; Doty Scientific Inc., Columbia, SC, USA) was placed above the skull, directly over the tumor. A ^13^C slice-selective, pulse-acquired spectroscopy sequence was prepared in which a single slice was placed over the tumor (FA = 25°, TE = 205 ms, TR = 2,000 ms, BW = 5 kHz, Matrix = 2048 x 64, FOV = 35 mm x 35 mm, NEX = 1, Slice Thickness = 6 mm, reference frequency = 75.515 MHz). Spectra were acquired every 2 seconds for 2 minutes to detect [1-^13^C]pyruvate and its lactate product.

#### Pyruvate-to-Lactate Measurements

The time-resolved stack of ^13^C MR spectra was imported into MATLAB (The MathWorks, Inc. Natick, MA, USA). A freely-available MATLAB script courtesy of the Hyperpolarized MRI Technology Resource Center (Hyperpolarized-MRI-Toolbox. Available online at: https://github.com/LarsonLab/hyperpolarized-mri-toolbox DOI: 10.5281/zenodo.1198915) was used to analyze the spectra. This script was adapted so that each of the individual spectra in the time-resolved stack could be phase- and baseline-corrected individually, rather than as a sum. Following these corrections, the pyruvate and lactate peaks were integrated (full-width, quarter-max), and the integral values were summed across all spectra in the time-resolved stack. The metric for pyruvate-to-lactate conversion, denoted as nLac, was then calculated using these values as the ratio of lactate:lactate+pyruvate. At least 5 nLac measurements from tumor-bearing mice were averaged at each time-point. Additionally, N = 3 nLac measurements from control mice were averaged and compared with tumor-bearing mice at these time-points.

#### Brain Sample Excision

Mice were euthanized and the bolt and top of the skull were carefully removed to expose the brain. The optical tracts were severed, and the intact brain was excised. For samples to be processed for immunohistochemistry (IHC) analysis, the entire brain was placed and fixed in a vial containing 10% formalin.

For samples to be processed for nuclear magnetic resonance (NMR) spectroscopy, incisions were made along the longitudinal and transverse fissures. The tissue within the left cerebral hemisphere (which contained the GSC injection site) was isolated from the remainder of the brain (which included the olfactory bulb, right cerebral hemisphere, and cerebellum). The isolated tissue was flash-frozen in liquid nitrogen and transferred to a freezer at -80°C. This process was executed as quickly as possible to preserve cellular metabolism, and the same volume of tissue was collected across all time-points, regardless of tumor size.

#### Sample Preparation for Metabolite Extraction

Metabolites were extracted from *ex vivo* samples as described in the literature (Weygand et al., 2017). Frozen samples were pulverized and weighed. The tissue particles were transferred to a conical centrifuge tube containing 2 mL of methanol, 1 mL of water, and 0.5 mL of lysing beads. The samples were vortexed, flash-frozen, and thawed for 3 cycles to lyse the cells. The samples were then centrifuged at 4°C for 10 minutes, and the supernatant, containing water, methanol, and dissolved metabolites, was extracted. The methanol was evaporated under reduced pressure using a rotary evaporator, and the remaining solvent was freeze-dried using a lyophilizer. The dried product was dissolved in 600 µL of deuterium oxide, 36 µL of phosphate buffered saline, and 4 µL of 4,4-dimethyl-4-silapentane-1-sulfonic acid (DSS-d6) as the NMR reference standard (500 µM in solution). The final 640 µL solution was transferred to a 5 mm NMR tube. All supplies (deuterium oxide, DSS, phosphate buffer) were purchased from Isotec Sigma Aldrich and used without further purification.

#### Nuclear Magnetic Resonance Spectroscopy

NMR spectra were obtained using a Bruker AVANCE III HD® NMR scanner (Bruker Biospin MRI GmbH, Ettingen, Germany) at a temperature of 298 K. The spectrometer operates at a ^1^H resonance frequency of 500 MHz and is endowed with a triple resonance (^1^H, ^13^C, ^15^N) Prodigy BBO cryogenic temperature probe with a Z-axis shielded gradient for increased sensitivity. A pre-saturation technique was implemented for water suppression. The spectra were obtained with a 90° pulse width, a scan delay t_rel_ of 6 s, a 10240 Hz spectral width, and an acquisition time t_max_ of 1.09 s (16,000 complex points). A total of 256 scans are collected and averaged for each spectrum, which results in a total scan time of 33 minutes. Here, t_rel_ + t_max_ is nearly 8 s so that it exceeds 3*T1 for the metabolites observed. The time domain signal is apodized using an exponential function.

#### Metabolite Pool Size Measurements

The raw FID files from the spectrometer were imported into MestreNova software (Mestrelab Research, A Coruña, Spain) where they were Fourier transformed into chemical shift spectra. The spectra were manually phase-corrected to form Lorentzian peak shapes and manually baseline-corrected to remove the noise floor, and the DSS reference peak was set to 0 ppm. The processed spectra were exported to Chenomx NMR Suite 8.1 software (Chenomx Inc., Edmonton, Canada) where the peaks were identified by matching them to spectral models of metabolites contained in the database. The identified peaks were integrated in MestreNova, and the peak areas were normalized to the peak area of DSS. As described in the literature (Zacharias et al., 2017), these values were further normalized to the mass of the pulverized tissue (in mg) and converted to metabolite concentrations (in µM) by implementing the Beer-Lambert law. The final values were reported as µM/mg. Similar to the nLac measurements, there were N = 5 groups of measurements for the tumors and N = 3 groups of measurements for the healthy brain controls at each time-point.

#### Sample Preparation for Immunohistochemistry

Mice were sacrificed by intracardiac perfusion of PBS and 4% paraformaldehyde. Brains were removed, fixed in 10% formalin for at least 24 hours and embedded in paraffin. Sections (5 µm) were cut for immunohistochemical analysis.

#### Immunohistochemistry

Sections of formalin-fixed paraffin-embedded mouse brain specimens were deparaffinized with xylene and rehydrated through a graded alcohol series, followed by distilled water and PBS. The slides were processed for antigen retrieval by pressure cooker in citrate buffer (pH 6.0) for 20 minutes. The slides were incubated overnight with the mouse antihuman MCT1 antibody (Santa Cruz Biotechnology, Dallas, TX, USA) at 1:200 dilution and rabbit antihuman LDH-A antibody (Abcam, Cambrige, MA, USA) at 1:100 dilution. For the MCT1 immunostaining, slides were incubated with Mouse Ig Blocking Reagent (Vector Laboratories, Burlingame, CA, USA) to reduce endogenous mouse Ig staining. The slides were rinsed with PBS and incubated with Anti-rabbit Poly-HRP-IgG (Leica Biosystems, Buffalo Grove, IL, USA) and Polymer Anti-Mouse IgG reagent (Leica Biosystems, Buffalo Grove, IL, USA) and visualized with 3,3′-Diaminobenzidine (DAB). The slides were counterstained with hematoxylin, dehydrated and coverslipped.

#### Protein Expression Measurements

The 40x images (1360 x 1024 pixels) of MCT1 and LDH-A stains were loaded into the FIJI package of ImageJ (Schindelin et al., 2012). Color deconvolution was performed (Ruifrok and Johnston, 2001) to extract the DAB stain from the image. These DAB images were thresholded so that DAB signal was set to 1 and everything else was set to 0. Three nonoverlapping, uniformly-sized ROI (340 x 256 pixels) were randomly placed in the tumor and one ROI outside the tumor as a background measurement. The percent area of DAB stain was calculated in these ROI (number of DAB-positive pixels divided by number of pixels in ROI). Percent area from the background ROI was subtracted from each of the tumor ROI which were then averaged together to produce the final metric for protein expression in that image. Average percent area from each image at a time-point were further averaged together to describe protein expression over the course of tumor development.

### QUANTIFICATION AND STATISTICAL ANALYSIS

All statistical analysis was conducted using GraphPad Prism 8 (GraphPad Software, La Jolla, CA, USA). All measurements are reported as the mean value ± standard deviation, and error bars in the figures represent standard deviation. N represents the sample size for each group as a single number or range of values.

#### Median Survival Time

Kaplan-Meier analysis was used to compare the median survival of untreated (N = 102) and treated (N = 59) tumor-bearing mice. Specifically, the Mantel-Cox logrank test was used to test for significant differences between median survival times, and the logrank method was used to calculate the hazard ratio.

#### In Vivo Tumor Volume Measurements

In untreated tumors during development, average volume at each time-point (N = 5-10) was compared with baseline average volume measured on Day 5 (N = 6). In treated tumors following radiotherapy, average volume at each time-point (N = 5-10) was compared with average volume at time of treatment on Day 26 (N = 8) as well as to maximum average volume (N = 10). Volume data was log-transformed to correct for heteroscedasticity and tested for significant differences using ordinary one-way ANOVA and follow-up Fisher’s Least Significant Differences tests (p < 0.05) where p is the probability value.

Repeated measures of individual tumor volume values in treated tumor-bearing mice were further analyzed using mixed-effects analysis with the Geisser-Greenhouse correction as an additional check for significant changes.

#### In Vivo Pyruvate-to-Lactate Measurements

During tumor development, average nLac of untreated tumors (N = 5) were compared with average nLac of controls (N = 3) at each time-point. In treated tumors, average nLac at each time-point (N = 5-8) was compared with average nLac at all prior time-points as well as to average nLac of untreated tumors and controls on Days 28 and 34. Statistical significance was determined using ordinary one-way ANOVA and follow-up Fisher’s Least Significant Difference tests (p < 0.05).

Repeated measures of individual nLac values in treated tumor-bearing mice were further analyzed using mixed-effects analysis with the Geisser-Greenhouse correction as an additional check for significant changes.

#### Ex Vivo Metabolite Pool Size Measurements

During tumor development, average metabolite pool sizes of untreated tumors (N = 5-7) were compared with average metabolite pool sizes of normal brain tissue controls (N = 3) at each time-point. In treated tumors, average metabolite pool sizes at each time-point (N = 5) were compared with average metabolite pool sizes at all prior time-points as well as to average metabolite pool sizes of untreated tumors and controls on Days 28 and 34. Statistical significance was determined using ordinary one-way ANOVA and follow-up Fisher’s Least Significant Difference tests. Due to the large number of comparisons (each set of comparisons made for 26 metabolites), the false discovery rate was controlled using the two-stage step-up method of Benjamini, Krieger and Yekutieli (Q < 0.05).

#### Ex Vivo Protein Expression Measurements

During tumor development, average background-subtracted percent stained area (MCT1 and LDH-A) from 3 ROI in each 40x capture (N = 1-5) from untreated tumors was compared at each time-point as well as with healthy brain controls. Statistical significance was determined using ordinary one-way ANOVA and follow-up Fisher’s Least Significant Difference tests (p < 0.05).

## Results

### Radiotherapy significantly extends survival of GSC 8-11 tumor-bearing mice

In mice implanted with GSC 8-11, a patient-derived GBM cell line, survival time was compared between untreated mice and mice treated with 2x5 Gy of whole-brain irradiation on Days 25 and 27 using Kaplan-Meier analysis (Fig. 2). Median survival was significantly increased in treated tumor-bearing mice compared with untreated mice (88 vs. 34 days, p < 0.0001). There was over a 250% increase in median survival time, which produced a hazard ratio of 4.6. Therefore, the radiotherapy dose of 2x5 Gy was effective at extending the survival of tumor-bearing mice, allowing for tumor regression and recurrence to be assessed in the radiotherapy treated mouse cohort.

**Figure 2:**
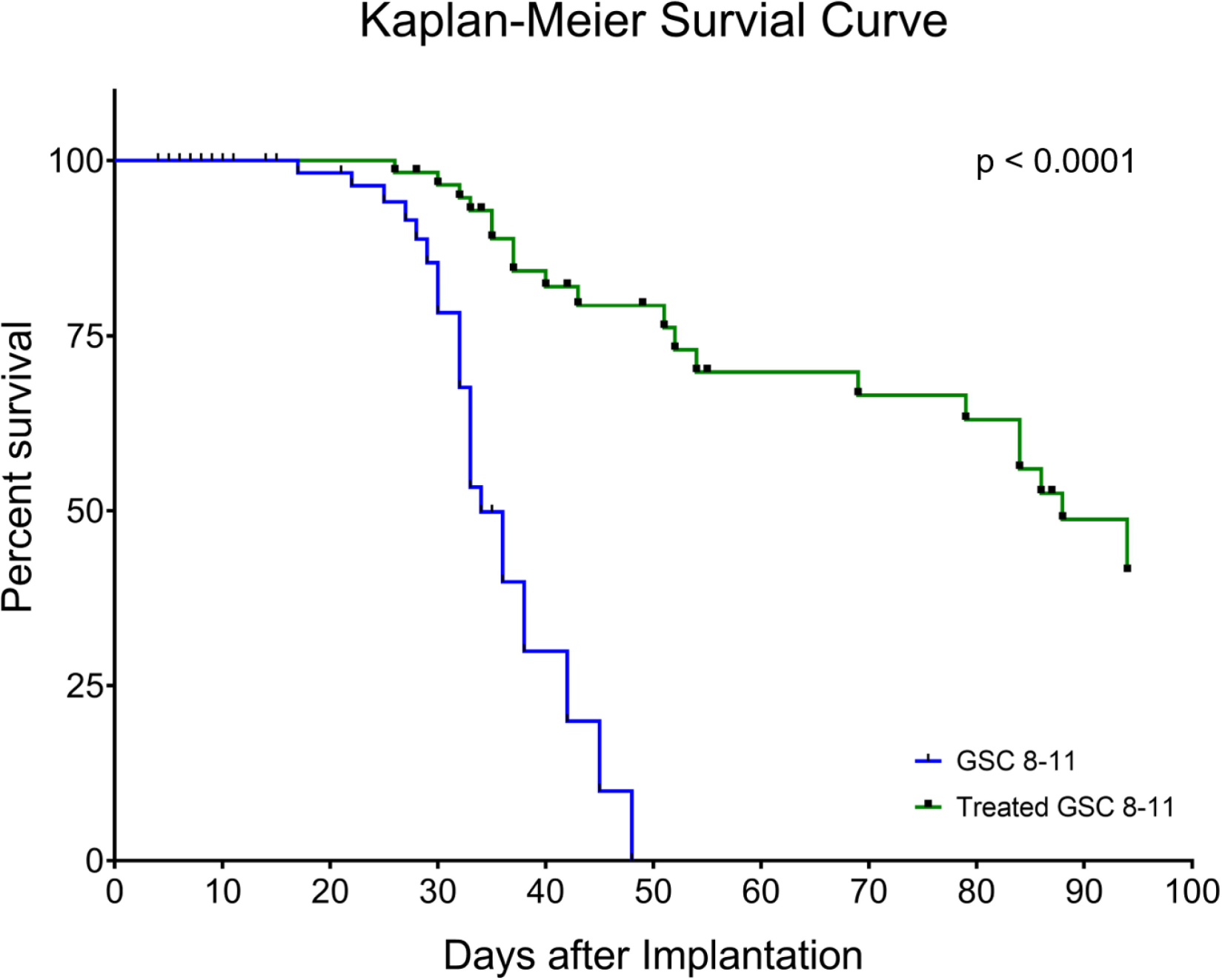
Radiotherapy significantly extends survival of GSC 8-11 tumor-bearing mice. Median survival of treated mice (green line) was significantly higher compared with untreated mice (blue line) (88 vs. 34 days, p < 0.0001). The hazard ratio of untreated-to-treated mice was 4.6. Mice that were euthanized on specific time-points for *ex vivo* experiments as well as those that died from non-tumor-related causes were censored (black markers). Survival was calculated using Kaplan-Meier analysis. Significant differences between the curves was calculated with the Mantel-Cox logrank test and the hazard ratio using logrank approach.

### Tumor volume increases during development but does not significantly change throughout regression or recurrence

#### Tumor Development (Day 1-34)

Following GSC implantation, average tumor volume was assessed every 3-4 days with T1-weighted, T2-weighted, and fluid-attenuated MRI (Fig. 3). An initial baseline volume of 1.4 ± 0.5 mm^3^ was measured on Day 5. Average tumor volume experienced exponential growth (R^2^ = 0.60), increasing slowly at first before rapidly expanding on Day 21 where it nearly tripled from the prior time-point to a value of 15.2 ± 9.7 mm^3^. At a value of 3.5 ± 0.9 mm^3^, average tumor volume was significantly increased by Day 10 compared with baseline volume (p = 0.0282). By the endpoint of tumor development (Day 34), average untreated tumor volume was 88.6 ± 56.3 mm^3^.

**Figure 3:**
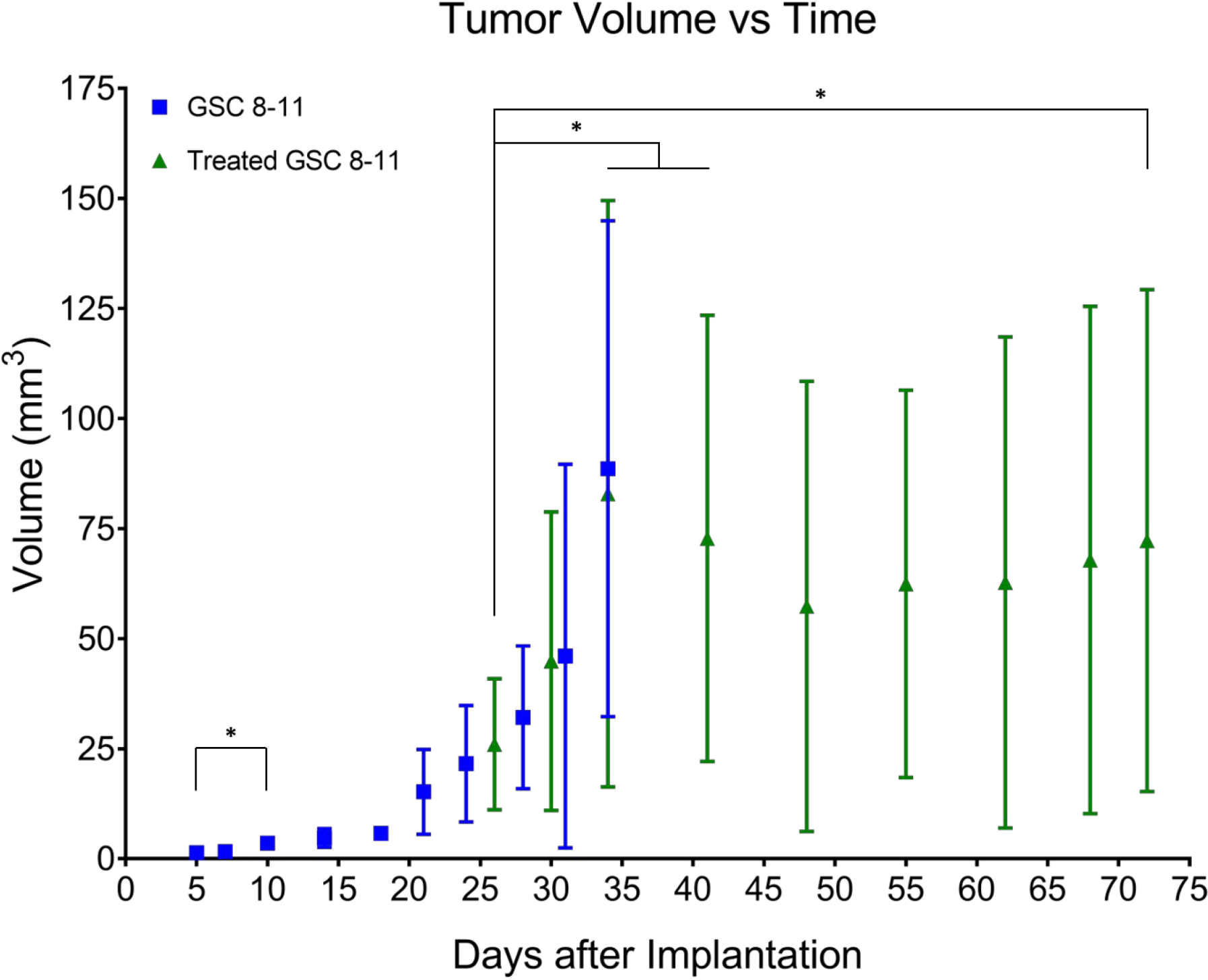
Tumor volume increases during development but does not significantly change throughout regression or recurrence. Average tumor volume, measured from anatomic MRI, is plotted as a function of time in untreated (blue squares) and treated (green triangles) mice. Error bars represent standard deviation (not visible before Day 21 due to scale of Y-axis). These values were log-transformed to correct for heteroscedasticity and tested for significant differences using ordinary one-way ANOVA and follow-up Fisher’s Least Significant Differences tests. Comparisons that produced p < 0.05 were deemed significant. *p < 0.05.

#### Tumor Regression (Day 25-48)

Following radiotherapy, tumor volume was measured on Days 26, 30, 34, and every 7 days thereafter (Fig. 3). On Day 34, average treated tumor volume was significantly increased compared with its initial value on Day 26 (82.9 ± 66.6 vs. 26.0 ± 14.9 mm^3^, p = 0.0090). Average treated tumor volume then began to decrease to a minimum of 57.3 ± 51.2 mm^3^ on Day 48 although no significant differences were observed compared with initial treated tumor volume on Day 26 or maximum treated tumor volume on Day 34.

#### Tumor Recurrence (Day 48-72)

The same group of mice were imaged every 7 days throughout tumor recurrence (Fig. 3). After reaching a minimum of 57.3 ± 51.2 mm^3^ on Day 48, average treated tumor volume monotonically increased to a final volume of 72.3 ± 57.0 mm^3^ on Day 72. While this value was significantly higher than initial treated tumor volume on Day 26 (p = 0.0396), neither grouped ANOVA analysis nor mixed-effects analysis of individual repeated measures revealed any significant increases in treated tumor volume throughout tumor recurrence.

### *In vivo* pyruvate-to-lactate conversion is significantly altered throughout tumor development, regression, and recurrence

#### Tumor Development (Day 1-34)

Using hyperpolarized ^13^C MRS, pyruvate-to-lactate conversion, quantified by nLac, was measured in tumor-bearing mice and control mice every 7 days with an extra measurement on Day 10 to capture early tumor dynamics (Fig. 4A). nLac is defined as the ratio of lactate:lactate+pyruvate where lactate and pyruvate are the area-under-the-curve values calculated from the time-resolved ^13^C spectra. Starting on Day 14, average nLac was significantly increased in tumor-bearing mice compared with control mice (0.39 ± 0.16 vs. 0.26 ± 0.10, p = 0.0273) which persisted throughout the remaining time-points in tumor development. In tumor-bearing mice, average nLac increased with a significantly nonzero slope (0.0069 days^-1^, p = 0.0006) to a final value of 0.45 ± 0.09 on Day 34. In contrast, average nLac in control mice remained constant across all time-points (slope = 0.0005 days^-1^, p = 0.7674), at an average value of 0.26 ± 0.07.

**Figure 4:**
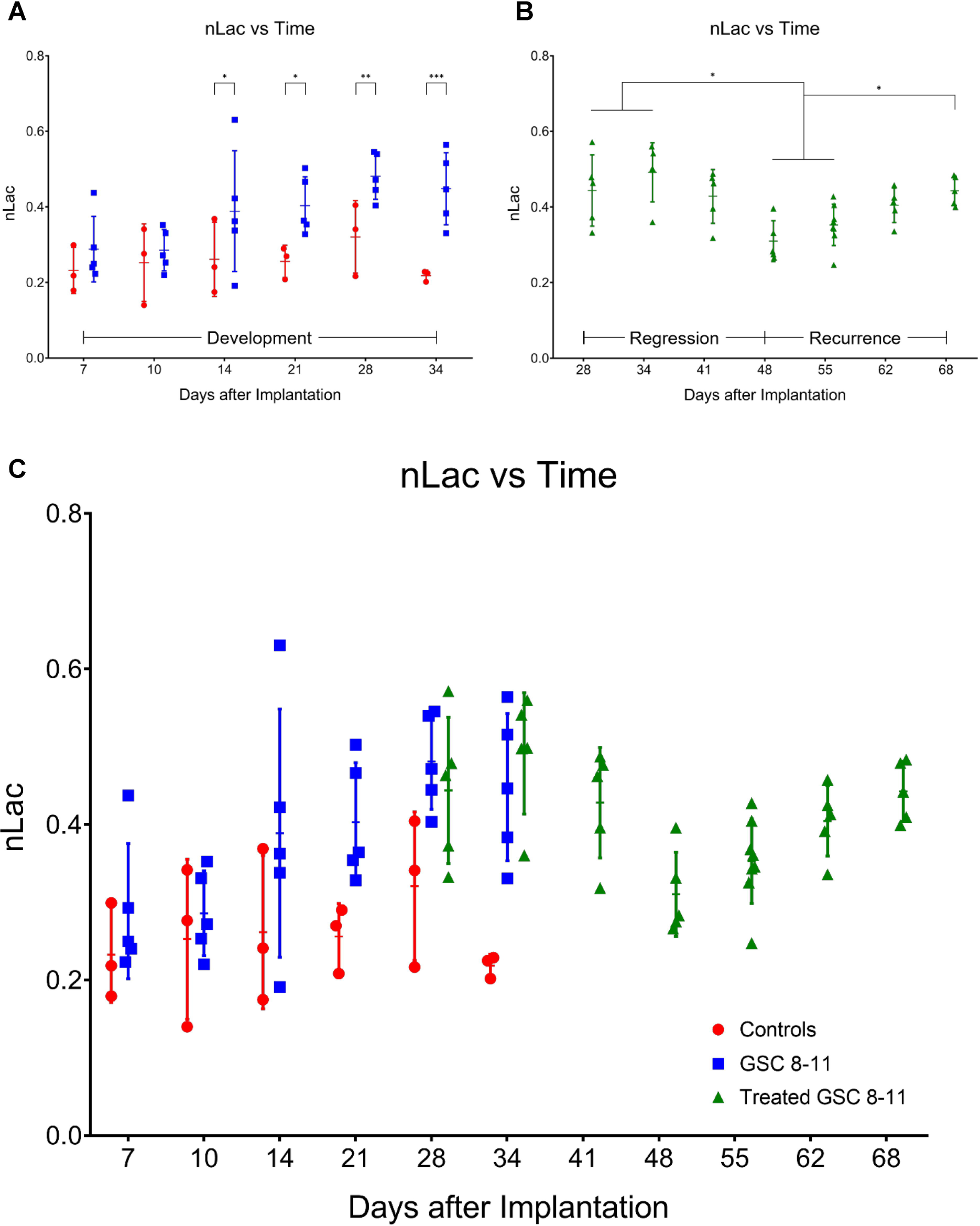
*In vivo* pyruvate-to-lactate conversion is significantly altered throughout tumor development, regression, and recurrence. Individual nLac values, measured with hyperpolarized ^13^C MRS, are plotted as a function of time for control mice (red circles) and untreated tumor-bearing mice (blue squares) during tumor development **(A)**. Individual nLac values are plotted as a function of time for treated tumor-bearing mice (green triangles) during tumor regression and recurrence **(B)**. Individual nLac values are plotted as a function of time across the entirety of tumor progression **(C)**. Error bars represent standard deviation. Average nLac values between groups and time-points were assessed for significance using ordinary one-way ANOVA and follow-up Fisher’s Least Significant Difference tests with significance attributed to comparisons that produced p < 0.05. *p < 0.05, **p < 0.01, ***p < 0.001.

#### Tumor Regression (Day 25-48)

Following radiotherapy, hyperpolarized MRS experiments were conducted in treated mice starting on Day 28 and every 7 days thereafter (Fig. 4B). Average nLac decreased following radiotherapy and was significantly decreased in treated tumor-bearing mice on Day 48 compared with treated tumor-bearing mice on Days 28 (0.31 ± 0.05 vs. 0.44 ± 0.09, p = 0.0080), 34 (0.31 ± 0.05 vs. 0.49 ± 0.08, p = 0.0004), and 41 (0.31 ± 0.05 vs. 0.43 ± 0.07, p = 0.0186). Additionally, average nLac was significantly decreased in treated tumor-bearing mice on Day 48 compared with untreated tumor-bearing mice on Days 28 (0.31 ± 0.05 vs. 0.48 ± 0.06, p = 0.0008) and 34 (0.31 ± 0.05 vs. 0.45 ± 0.09, p = 0.0063). Lastly, of the tumor-bearing mice which had initial values of nLac < 0.4 immediately following treatment (on Day 28 or 34), 0/3 died from tumor burden by the Day 94 endpoint of the study. In contrast, 5/7 treated tumor-bearing mice with initial values of nLac > 0.4 died from tumor burden before reaching the Day 94 endpoint.

#### Tumor Recurrence (Day 48-72)

Hyperpolarized MRS experiments were performed on the same group of mice every 7 days throughout tumor recurrence (Fig. 4B). Average nLac linearly increased with a significantly nonzero slope (0.0067 days^-1^, p < 0.0001) in treated tumor-bearing mice from Day 48 to Day 68. When compared with the trend of average nLac in untreated tumor-bearing mice during tumor development, average nLac in treated tumor-bearing mice during tumor recurrence had a nearly identical slope (0.0067 vs. 0.0069 days^-1^, p = 0.9415). When analyzed with grouped ANOVA analysis, average nLac in treated tumor-bearing mice was significantly increased by Day 68 compared with treated tumor-bearing mice on Days 48 (0.44 ± 0.04 vs. 0.31 ± 0.05, p = 0.0085) and 55 (0.44 ± 0.04 vs. 0.35 ± 0.05, p = 0.0452) as well as when compared with control mice on Days 28 (0.44 ± 0.04 vs. 0.32 ± 0.10, p = 0.0341) and 34 (0.44 ± 0.04 vs. 0.22 ± 0.01, p = 0.0002). Furthermore, when analyzing comparisons of individual repeated measurements with mixed-effects analysis, nLac was still significantly increased on Day 68 compared with Day 55 (0.44 ± 0.04 vs. 0.35 ± 0.05, p = 0.0214).

### *Ex vivo* metabolite pool sizes are significantly altered throughout tumor development and regression

#### Tumor Development (Day 1-34)

Tumors and controls (healthy murine brain tissue) were excised at the same time-points as the hyperpolarized MRS experiments for *ex vivo* global metabolomics using NMR spectroscopy. Pool sizes of 26 metabolites were quantified and compared between tumors and controls in the same manner as nLac (Fig. 5). By Day 28, average pool size of the metabolites alanine (3.92 ± 1.55 vs. 1.54 ± 0.61 µM/mg, q = 0.0366) and phosphocholine (3.14 ± 1.62 vs. 1.32 ± 0.49 µM/mg, q = 0.0491) were significantly increased in tumors compared with controls. On Day 34, average pool sizes of alanine (4.80 ± 1.98 vs. 1.45 ± 0.97 µM/mg, q = 0.0027) and phosphocholine (3.40 ± 1.23 vs. 1.17 ± 0.64 µM/mg, q = 0.0144) were still significantly increased in tumors compared with controls along with glycerophosphocholine (3.19 ± 1.14 vs. 0.93 ± 0.34 µM/mg, q = 0.0343), glycine (10.56 ± 6.53 vs. 2.86 ± 2.01 µM/mg, q = 0.0106), and valine (0.81 ± 0.43 vs. 0.28 ± 0.20 µM/mg, q = 0.0072).

**Figure 5:**
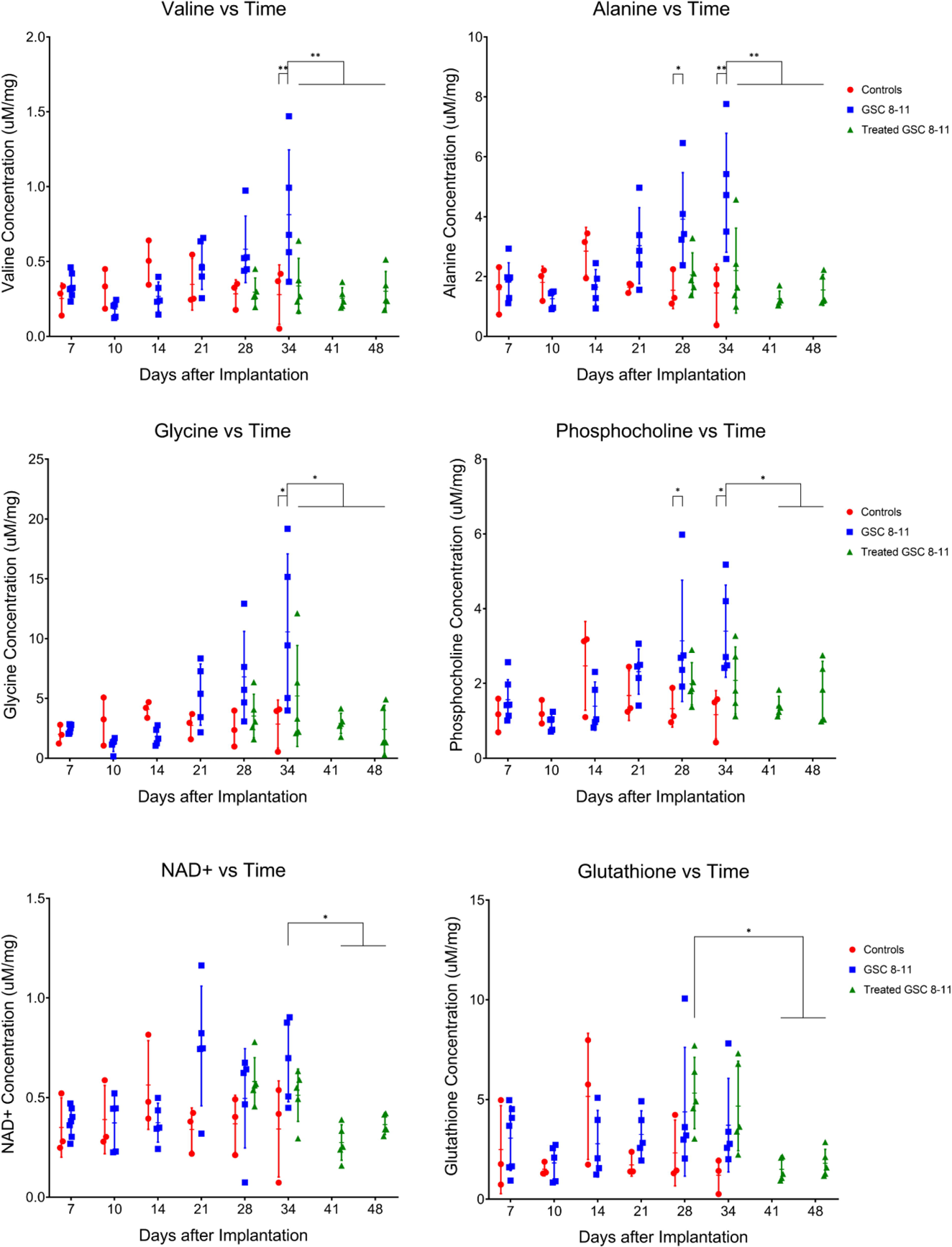
*Ex vivo* metabolite pool sizes are significantly altered throughout tumor development and regression. Individual metabolite pool sizes, measured with NMR spectroscopy, are plotted as a function of time for control mice (red circles) and untreated (blue squares) and treated (green triangles) tumor-bearing mice. Error bars represent standard deviation. The Y-axis is reported as concentration of metabolite pool per mg of tissue. Statistical significance was determined using ordinary one-way ANOVA and follow-up Fisher’s Least Significant Difference tests. The false discovery rate was controlled using the two-stage step-up method of Benjamini, Krieger and Yekutieli, and significance was attributed to comparisons that produced q < 0.05. *q < 0.05, **q < 0.01.

#### Tumor Regression (Day 25-48)

Following radiotherapy, samples were excised from treated mice at the same time-points as the hyperpolarized MRS experiments for NMR spectroscopy (Fig. 5). On Day 34, average pool size of the following metabolites were significantly decreased in treated tumors compared with untreated tumors: alanine (2.20 ± 1.42 vs. 4.80 ± 1.98 µM/mg, q = 0.0072), glycine (5.21 ± 4.23 vs. 10.56 ± 6.53 µM/mg, q = 0.0457), and valine (0.34 ± 0.18 vs. 0.81 ± 0.43 µM/mg, q = 0.0061). By Day 48, average pool size of these metabolites were still significantly decreased in treated tumors compared with untreated tumors on Day 34 along with the metabolites, NAD+ (0.36 ± 0.05 vs. 0.69 ± 0.21 µM/mg, q = 0.0496) and phosphocholine (1.80 ± 0.79 vs. 3.40 ± 1.23 µM/mg, q = 0.0457). Another result of note is that on Day 28, average glutathione pool size was significantly increased in treated tumors compared with treated tumors on Days 41 (5.32 ± 1.79 vs. 1.50 ± 0.58 µM/mg, q = 0.0328) and 48 (5.32 ± 1.79 vs. 1.81 ± 0.70 µM/mg, q = 0.0491). Refer to Table S1 for all significant metabolite pool size changes throughout tumor development and regression.

### *Ex vivo* MCT1 expression significantly increases throughout tumor development

Immunohistochemistry (IHC) was performed on *ex vivo* tumor samples at several time-points to measure changes in MCT1 and LDH-A expression throughout tumor development (Fig. 6). Qualitatively, MCT1 was membrane-bound and LDH-A was confined to the cytoplasm, as expected. Semi-quantitatively, average MCT1 percent stained area was significantly higher in untreated tumors by Day 21 compared with controls (13.43 ± 5.33 vs. 3.11 ± 3.57, p = 0.0139; 13.43 ± 5.33 vs. 2.73 ± 1.98, p = 0.0110) and remained significantly increased through Days 25, 28, and 34. Additionally, average MCT1 percent stained area was significantly higher in untreated tumors by Day 21 compared with untreated tumors early in development on Day 10, (13.43 ± 5.33 vs. 5.15 ± 4.86, p = 0.0459) and remained significantly increased through Days 25, 28, and 34. Conversely, average LDH-A percent stained area was elevated in untreated tumors at all time-points of development, and the only significant increase was on Day 34 when compared with Days 14 (37.15 ± 5.48 vs. 19.77 ± 6.88, p = 0.0002), 25 (37.15 ± 5.48 vs. 23.86 ± 5.52, p =0.0030), and 28 (37.15 ± 5.48 vs. 20.24 ± 12.81, p = 0.0008).

**Figure 6:**
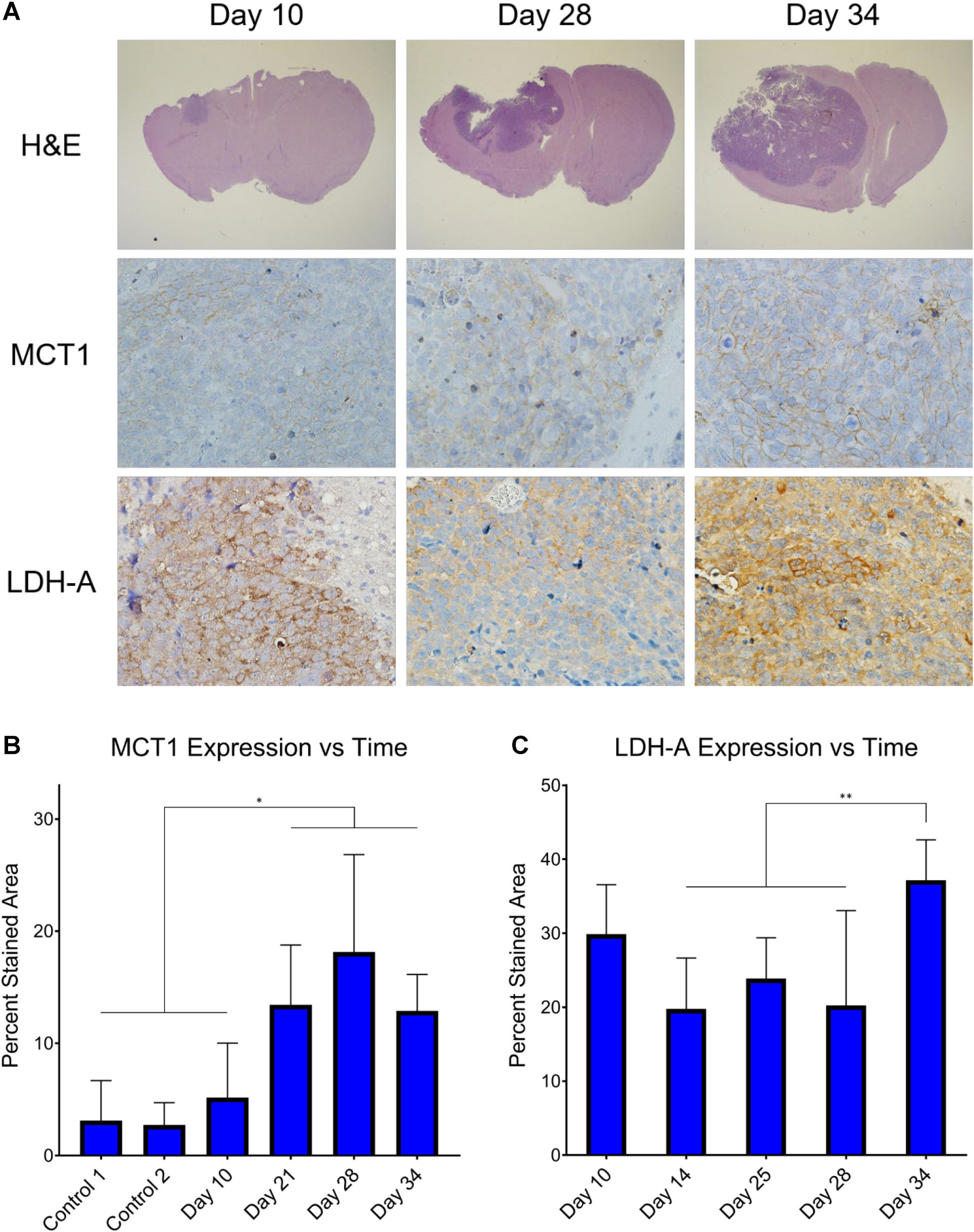
*Ex vivo* MCT1 expression significantly increases throughout tumor development. Histology stains of H&E, MCT1, and LDH-A were measured in *ex vivo* untreated tumor samples at several time-points throughout tumor development **(A)**. It can be clearly seen that MCT1 is confined to the cell membrane and LDH-A to the cytoplasm. Percent stained area was calculated from the MCT1 **(B)** and LHD-A **(C)** IHC images at these time-points. Average percent stained area between time-points was assessed for significance using ordinary one-way ANOVA and follow-up Fisher’s Least Significant Difference tests with significance attributed to comparisons that produced p < 0.05. *p < 0.05, **p < 0.01.

## Discussion

Our findings demonstrate that *in situ* analysis of metabolic changes linking the stages of tumor progression (tumor development, regression following radiotherapy, and recurrence) using hyperpolarized MRI and NMR spectroscopy is feasible and informative. By acquiring each of these measurements, along with anatomic growth, in the same mice and at multiple common time-points, the individual evolution of the results from these assays were investigated as well as their relationship and correlation with one another. Thus, an extensive evaluation of cancer metabolism as it advances through different stages of tumor progression was examined. The results from these experiments illustrate that measured pyruvate-to-lactate conversion values are variable in all stages of tumor progression. Therefore, when researchers are using this technique to compare hyperpolarized pyruvate-to-lactate values between tumor types of varying growth rates or before and after treatment, the stage of tumor progression needs to be considered. The mouse model used in this project is an orthotopic implantation of patient-derived GBM (GSC 8-11), which generates confidence that these results are clinically relevant and potentially translatable. However, we do acknowledge the caveat that host immunity is not fully represented in these models.

To our knowledge, this is the first study to investigate all three stages of brain tumor progression, including tumor recurrence, with hyperpolarized MRS. There is one prior study that used hyperpolarized MRSI to measure the glycolytic effects of switching off MYC expression in transgenic mouse models of breast cancer which resulted in tumor regression followed by recurrence (33). Shin, et al. observed that the hyperpolarized lactate:pyruvate ratio significantly decreased as the tumors regressed from MYC withdrawal, and after a latency period, the tumors recurred, which was accompanied by a significant increase of hyperpolarized lactate:pyruvate ratio. In one of the mice, this increase of lactate production was observed prior to increases in tumor volume, which supports the results we observed across multiple treated GBM tumor-bearing mice.

Because average tumor volume values and their variances were low on Day 5, significant increases in untreated tumor volume were detected as soon as Day 10. However, it was not until Day 21 that average tumor volume began to rapidly increase, tripling in value compared with Day 18. Meanwhile, nLac was significantly increased compared with controls beginning on Day 14. Thus, hyperpolarized MRS predicted aggressive growth while the tumor was still in the slow-growing phase. One potential clinical application of this is the ability to distinguish aggressive tumors from their slower-growing counterparts early in tumor development and predict malignant transformation. This would allow physicians to begin administering therapy before the tumor aggressively invades neighboring tissue and is still treatable, which may lead to improvements in patient survival.

Another current clinical challenge facing neuro-oncologists is distinguishing pseudoprogression from true progression following radiotherapy. Pseudoprogression is an anatomic MRI pattern that mimics tumor progression and can confound treatment monitoring with direct consequences in clinical practice. It can lead to prematurely withholding adjuvant temozolomide or continuing with potentially ineffective treatment in patients in which cases of tumor progression are not clear (34). Pseudoprogression was observed in this study with anatomic MRI as tumor volumes in treated mice significantly increased on Days 34 and 41 before eventually decreasing throughout regression. Conversely, there were no significant increases in nLac during this time-course, which suggests hyperpolarized MR could help mitigate this problem in the clinic.

Furthermore, the predictive value of hyperpolarized MRS to stratify subjects into likely and unlikely responders was alluded to when looking at the initial nLac values following treatment. Of the 10 mice which underwent hyperpolarized MRS immediately following treatment, a cutoff value of nLac > 0.4 was able to predict if a mouse would succumb to tumor burden before the Day 94 endpoint with a sensitivity of 100% and specificity of 60%. These sample sizes are too small to draw definitive conclusions, but it serves as motivation for a larger study.

As the treated tumors proceeded through the regression stage following radiotherapy, nLac decreased significantly in treated tumor-bearing mice on Days 48 and 55 compared with Days 28 and 34, whereas tumor volume did not significantly decrease in this time period. When comparing the quality of the data, there were larger magnitudes of change and lower variance in the hyperpolarized MRS data compared with the anatomic MRI data. This remained true when looking at relative changes of nLac and tumor volume (by normalizing parameters to their initial value following therapy for those mice that were imaged several times throughout regression and recurrence). Within this set of repeated tumor volume measurements, average nLac fell to a minimum of 71% of its initial value after treatment by Day 55 whereas average tumor volume reached a minimum of 81% of its initial treated volume on Day 62 (Fig. 7). This suggests that hyperpolarized MRS can effectively detect the response of GBM to radiotherapy and do so more reliably than measuring changes in tumor volume with anatomic MRI.

**Figure 7:**
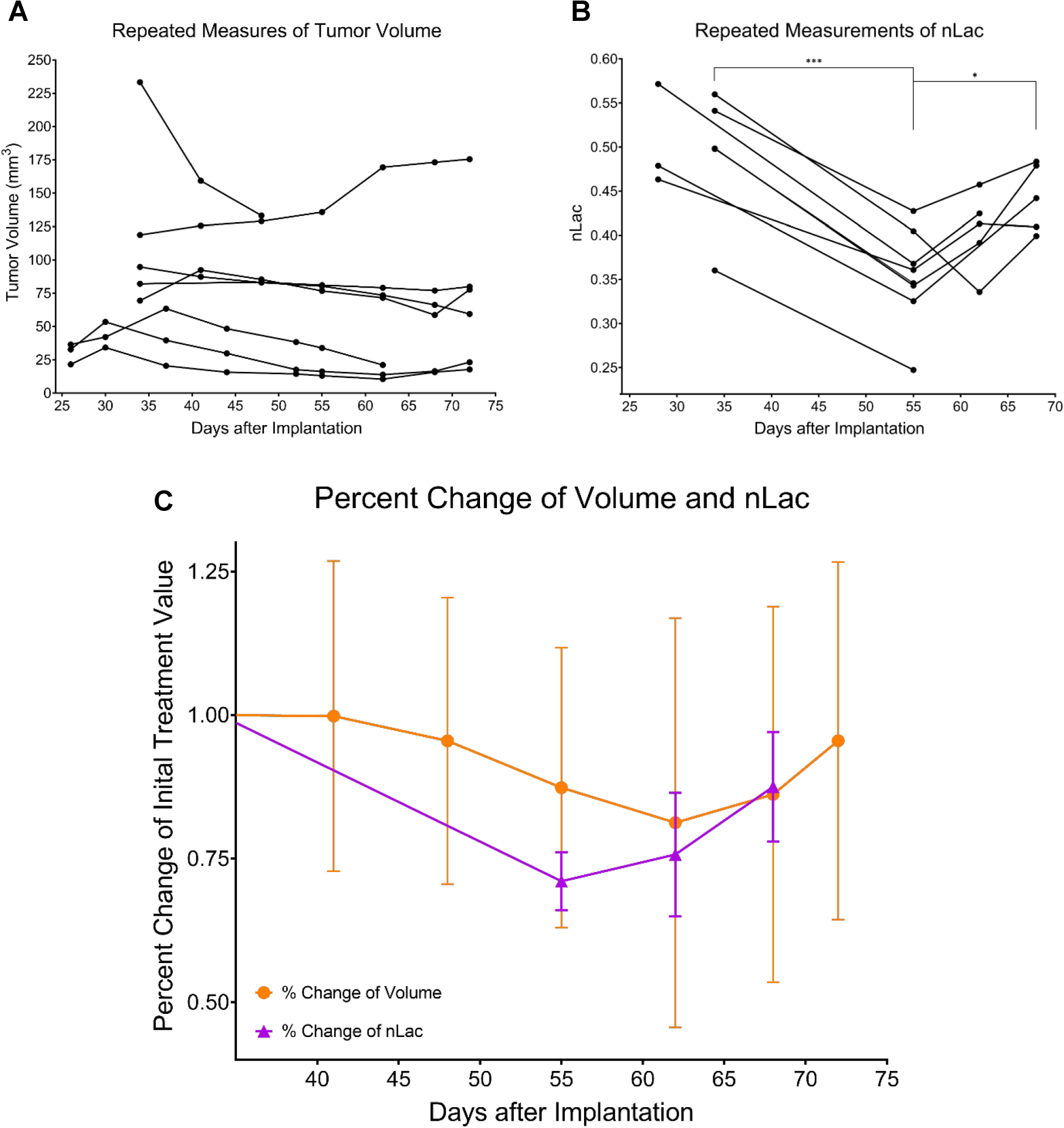
Percent change of nLac, but not tumor volume, is significantly altered during tumor regression and recurrence. Repeated measures of tumor volume are acquired over time in treated mice with anatomic MRI **(A)** and hyperpolarized MRS **(B)**. At each time-point, volume and nLac were normalized to their initial value following treatment and plotted as percent change over time **(C)**. The data were tested for significance using mixed-effects analysis with the Geisser-Greenhouse correction, and comparisons that produced p < 0.05 were deemed significant. *p < 0.05, ***p < 0.001.

Hyperpolarized MRS was also effective at predicting relapse as nLac was significantly increased in treated mice by Day 68 compared with Days 48 and 55 whereas tumor volume did not significantly increase in this time period. The only significant change in volume that was observed at any time-point following radiotherapy was that average treated tumor volume on Day 72 was significantly increased compared with initial treatment volume on Day 26, which does suggest that the mice were beginning to relapse. Furthermore, nLac values in treated mice on Day 68 were equivalent to initial values following treatment on Days 28 and 34. Thus, there was a complete reversal of nLac behavior by Day 68, further supporting that relapse was occurring. Again, when looking at percent change of nLac and tumor volume in the mice that were measured at multiple time-points throughout regression and recurrence (Fig. 7), average nLac significantly increased from its minimum of 71% of its initial value following treatment to 88% by Day 68 (p = 0.0258). Average tumor volume increased from its minimum of 81% of its initial value following treatment to 96% by Day 72 (p = 0.1084). Based on these data, we suspect that pyruvate-to-lactate conversion measured with hyperpolarized MRS in this animal model is a direct metabolic readout of tumor response during regression (Day 55) and relapse (Day 68) which is occurring prior to radiological tumor volume changes.

Another interesting result is that the slope of nLac in treated tumor-bearing mice in the tumor recurrence stage was nearly identical to that of the untreated tumor-bearing mice in the development stage. This suggests that the metabolic programming of recurring tumors was not altered, and the mechanism of recurrence was similar to that of initial growth. This is expected since tumors were only treated with radiotherapy. Had adjuvant chemotherapy or targeted therapy been implemented, cancer cells could have potentially been pressured to escape therapy through the development of resistance by adapting and reprogramming a new mechanism of survival and proliferation, which may have been reflected in the metabolic assays. This idea is currently of immense interest as metabolic changes that reflect resistance to therapy could be exploited with molecular imaging techniques and would be invaluable to physicians looking to optimize therapeutic approaches. This could have a significant benefit for patients with recurrent disease, and future research will investigate this idea further.

It has been traditionally accepted that an increase of the enzyme lactate dehydrogenase-A (LDH-A) accompanies tumor formation and is the driver of increased pyruvate-to-lactate conversion observed in hyperpolarized MR studies. However, the only significant increase of LDH-A expression, as measured with IHC, observed in this study was at the final time-point of tumor development on Day 34. *Ex vivo* lactate pool size in the GSC 8-11 tumors from NMR experiments did not reveal significant changes over the course of tumor progression. Additionally, neither untreated nor treated tumors produced significant correlations between their *in vivo* nLac values with *ex vivo* lactate pool sizes although we have observed this correlation in patient-derived pancreatic cancer mouse models of increasing aggressiveness (35). While the role of monocarboxylate transporter 1 (MCT1) for transporting pyruvate to the cytosol has been known to be important in hyperpolarized MR exams, it was recently demonstrated that MCT1 may be the key and rate-limiting step for lactate production (36). Additional studies report correlations of MCT1 with hyperpolarized pyruvate-to-lactate conversion in GBM (23) and breast cancer (37). Our data seems to support these results as MCT1 significantly increased throughout tumor development and mirrored the trajectory of nLac.

An analysis of the REMBRANDT and TCGA databases of patient brain tumors revealed that *SLC16A1* expression- the gene which encodes MCT1- was elevated in GBM compared with lower-grade astrocytoma, oligodendroma, and non-tumor brain samples (38). When grouping these glioma subtypes together, those with high expression of *SLC16A1* led to significantly lower survival compared to those with low expression of *SLC16A1* (39). Further analysis of the TCGA database revealed that MCT1 expression was significantly reduced in IDH1-mutant glioma samples compared to those with wild-type IDH1 (40). IDH1 mutations are predominately found in WHO grade II and III gliomas (70% of cases) compared to grade IV GBM (12% of cases), and GBM which harbor the IDH1 mutation lead to significantly longer survival times of greater than two-fold (41, 42). In individual studies, MCT1 immunoreactivity scores were significantly increased in 24 high-grade patient samples of GBM and anaplastic astrocytoma compared with 24 low-grade patient samples of oligodendrogliomas and low-grade astrocytomas (43), and it was significantly increased in 78 GBM patient samples compared with 24 non-tumor brain samples (44). Furthermore, the MCT inhibitor α-cyano-4-hydroxycinnamate (CHC) induced cytotoxic effects and inhibited proliferation, invasion, and migration capacity in high-grade glioma cell lines which possessed high expression of MCT1 (44, 45). Because MCT1 has differential expression between malignant and healthy brain cells, and even between low- and high-grade disease, it is an attractive prognostic biomarker. In this study, we have shown that increasing MCT1 expression correlates with hyperpolarized pyruvate-to-lactate conversion throughout GBM tumor development. In patients with prostate (46) and breast cancers (37), similar correlations of MCT1 expression and hyperpolarized pyruvate-to-lactate conversion were observed which also correlated with tumor grade. Thus, hyperpolarized MRS has a promising clinical application in the early detection of high-grade brain cancer through MCT1 interrogation.

In addition to probing changes in glycolytic metabolism, we were also interested in identifying alternate metabolic pathways which were potentially deregulated throughout tumor progression. These could serve as leads for subsequent imaging probes and therapeutic targets. Results from the NMR experiments of the *ex vivo* tissue samples demonstrated that amino acid metabolism was significantly altered throughout tumor progression. These metabolites are essential for protein synthesis and are necessary for cellular growth and division. In particular, the pool sizes of valine, alanine, and glycine were significantly increased by the end of tumor development and decreased following radiotherapy. Valine is a branched chain amino acid (BCAA) and can be used for protein synthesis or oxidized for energy production. Branched-chain aminotransferase 1 (BCAT1) generates glutamate during BCAA catabolism and is overexpressed in many cancers including glioma (47–50). Alanine and α-ketoglutarate can be reversibly produced from pyruvate and glutamate through alanine transaminase (ALT) whenever the pyruvate substrate is available. In some hyperpolarized [1-^13^C]pyruvate MR experiments, it is possible to observe hyperpolarized alanine production, and a decreasing ratio of hyperpolarized alanine-to-lactate has been suggested as a biomarker of disease progression in pancreatic cancer (51, 52). Unfortunately, we could not reliably observe hyperpolarized alanine in our experiments to suggest the same is true for GBM. Increased glutamine anaplerosis via ALT has been linked to the viability and proliferation of brain (53), breast (54, 55), colorectal (56), and prostate (57) cancers. It was recently demonstrated that alanine uptake and utilization through the SLC38A2 membrane transporter played a key role in pancreatic cancer metabolism and proliferation (58). Glycine is derived from serine in one-carbon metabolism to maintain redox balance through antioxidant production such as glutathione as well as to produce metabolites involved in purine nucleotide and lipid synthesis, all of which are important for cancer survival and proliferation (59). Glycine production through serine hydroxymethyltransferase (SHMT)- a transcriptional target of c-Myc (60)- has been implicated as a driver of cancer cell proliferation in glioma (61) and many other types of tumors (62–66).

Many tumors rely on antioxidants to quench the effects of reactive oxygen species (ROS) produced from treatments such as radiotherapy and chemotherapy (67) as well as oxidative stress from increased energy metabolism (68). This is often seen through an increased production of NADPH and glutathione through the pentose phosphate pathway and one-carbon metabolism (69). We observed a significant increase in *ex vivo* glutathione pool size in treated tumors one day following radiotherapy compared with treated tumors further into regression, which we believe is an acute response to increased ROS generated from radiotherapy. Increased concentration of antioxidants such as glutathione in tumors immediately following treatment has also been reported elsewhere (70, 71).

Increased phospholipid metabolism was observed in these GBM tumors compared with normal brain tissue and correlated with progression. The Kennedy pathway describes the phosphorylation of choline and ethanolamine to phosphocholine and phosphoethanolamine, which eventually form phosphatidylcholine and phosphatidylethanolamine (72). These are the two most abundant phospholipids in the cell membrane. The second messenger diacylglycerol is produced in this pathway which can further activate downstream signaling for cellular growth and fatty acid oxidation (73). Phosphatidylcholine can be broken down into glycerophosphocholine for storage and eventually converted back into choline. Increased phosphocholine and choline-containing metabolite concentrations have been observed in gliomas (53,74–77) and many other types of cancer (78–80), so it was not surprising to see significantly elevated pools of these metabolites, along with phosphoethanolamine in untreated tumors compared with controls and treated tumors.

Nicotinamide adenine dinucleotide (NAD+) is an important cofactor in cellular metabolism and is necessary for glycolysis, pyruvate-to-lactate conversion, and serine biosynthesis. Nicotinamide phosphoribosyltransferase (NAMPT) is the main enzyme for NAD+ biosynthesis and has been found to be upregulated in several cancers including glioma (81, 82). Inhibition of this enzyme leads to antitumoral effects, which has led to the development of drugs for different types of cancer (83–87). We observed a significant decrease in NAD+ levels following treatment.

In addition to uncovering potential diagnostic and therapeutic targets of metabolism, the data from this analysis can be combined with *in vivo* hyperpolarized MR data to build a model based on tumor metabolism to predict clinical outcomes. Correlative analysis demonstrated that hyperpolarized MR and NMR spectroscopy data are largely orthogonal (with hyperpolarized MR measuring real-time flux and NMR spectroscopy measuring metabolite pool size), so this model should be more predictive than either technique alone. Unfortunately, we could not implement this in our study because the tumor excision process for NMR spectroscopy requires euthanasia of the mice, preventing validation of treatment response or survival outcomes. However, in the clinical setting, biopsies of tumor tissue can be obtained during diagnosis as well as post-surgery, which could then be processed for NMR spectroscopy. Combining this data with hyperpolarized MR acquisitions at these time-points to predict clinical outcomes could form the basis for a clinical trial.

## Conclusion

This study demonstrated and discussed the benefits that hyperpolarized MRS could add to conventional clinical imaging to address several clinical challenges in the diagnosis and treatment of GBM. These include the ability to predict whether a tumor will be slow-growing or aggressive at the time of diagnosis, help discriminate pseudoprogression from true progression and predict whether patient survival will be improved shortly after administration of a treatment, and determine whether the patient is on the verge of relapse during a follow-up exam. Each of these scenarios would give physicians the time to take appropriate interventional action, improving the chances of patient survival.

*In vivo* metabolic measurements with hyperpolarized MRS were supported by *ex vivo* global metabolomics with NMR spectroscopy and protein expression assays with IHC to provide a comprehensive analysis of tumor metabolism. A major innovation in this study is that these measurements of metabolism, along with tumor volume, were made at several time-points across tumor development, regression, and recurrence. Thus, the individual evolution of tumor volume, hyperpolarized pyruvate-to-lactate conversion, and *ex vivo* metabolite pool sizes could be studied as well as their correlations with each other over time. As hyperpolarized MR makes its way through clinical trials as a metabolic imaging modality, we believe its value in cancer care will continue to grow. Just as positron emission tomography (PET) became a staple in the clinic, so too should hyperpolarized MRI as an invaluable tool for interrogating the metabolism of cancer.

## Acknowledgements

We gratefully acknowledge Mr. Jorge Delacerda, Mr. Charles Kingsley, Dr. James Bankson, and Dr. John Hazle for support with animal imaging. TCS also acknowledges Dr. Richard Wendt and Dr. Ho-Ling Anthony Liu for graduate advisory support. TCS further acknowledges Dr. Prasanta Dutta, Dr. Shivanand Pudakalakatti, and Dr. Jaehyuk Lee for experimental training.

