## Supplemental Table 1 for "Measuring the metabolic evolution of glioblastoma throughout tumor development, regression, and recurrence with hyperpolarized magnetic resonance"

| Metabolite | Tumor Development | Tumor Regression | Potential Pathway |
| --- | --- | --- | --- |
| Valine | U34 > C34, q=0.0072 | T34 < U34, q=0.0061  T41 < U34, q=0.0013  T48 < U34, q=0.0027 | BCAA Catabolism |
| Alanine | U28 > C28, q=0.0366  U34 > C34, q=0.0027 | T34 < U34, q=0.0072  T41 < U28, q=0.0063  T41 < U34, q=0.0004  T48 < U28, q=0.0149  T48 < U34, q=0.0011 | Glutamine Anaplerosis |
| Glycine | U34 > C34, q=0.0106 | T34 < U34, q=0.0457  T41 < U34, q=0.0034  T48 < U34, q=0.0021 | Glycine Cleavage  Folate Cycle |
| Phosphocholine | U28 > C28, q=0.0491  U34 > C34, q=0.0144 | T41 < U28, q=0.0284  T41 < U34, q=0.0106  T48 < U34, q=0.0457 | Kennedy Pathway  Choline Cycle |
| Glycero-phosphocholine | U34 > C34, q=0.0343 |  |  |
| Phosphoethanolamine |  | T41 < U28, q=0.0154  T41 < U34, q=0.0496  T48 < U28, q=0.0401 |  |
| Glutathione |  | T41 > T28, q=0.0328  T48 > T28, q=0.0491 | Trans-Sulphuration Pathway |
| NAD+ |  | T41 < U34, q=0.0106  T48 < U34, q=0.0496 | Energy Metabolism |

Table S1: List of significantly altered metabolites during tumor development and/or tumor regression. In the “Tumor Development” and “Tumor Regression” columns, the naming convention is the group (U=untreated, C=control, T=treated) followed by the time-point. For example, U34 > C34, q=0.0241 indicates that the metabolite was significantly increased in untreated tumor-bearing mice on Day 34 compared with control mice on Day 34, and the comparison produced a q-value of 0.0241. The q-values are p-values adjusted to control the false discovery rate using the two-stage step-up method of Benjamini, Krieger and Yekutieli. Pathways that the metabolites belong to which have been reported to be upregulated in cancer are also presented.
